# Design and Test Novel EEG Electrode Attachments for Type 4 Afro-textured Hairstyles

**DOI:** 10.64898/2026.10.01.755885

**Authors:** Mishti Broor, Kenneth Ogunniyi, Lucinda Williamson, Chang Liu

## Abstract

Black participants remain underrepresented in electroencephalography (EEG) research, partly because current EEG devices work poorly with dense, coily Type 4 Afro-textured hair. Research-grade EEG typically wires electrodes on a cap, and therefore the electrodes often fail to reach the scalp through dense, Afro-textured hair. Protocols that ask Black participants to unbraid or restyle protective hairstyles add burdens that discourage enrollment in research studies. To address this, we designed and tested 3D-printed attachments that fit onto an EEG electrode and anchored it directly in the hair, using the hairstyle itself as the mounting structure.

We fabricated three design configurations for natural Afro-textured hair, cornrows/braids, and twists/microlocs. Seventeen healthy adults with Type 4a-4c hair used electrode attachments that matched their hairstyle. During a single visit, EEG was recorded with 16 conventional cap electrodes defined by the international 10–20 system and with 16 electrodes using novel electrode attachments anchored directly to their hair. Participants completed resting and walking trials with each system. Electrode-to-scalp distance and gel volume were measured during setup. Electrode disconnection, impedance, signal quality (i.e. peak alpha power and aperiodic (1/f) component), and comfort level were measured throughout the data collection.

Compared with the conventional cap, the novel attachments reduced electrode-to-scalp distance, gel use, and impedance. Twenty percent of the conventional cap electrodes failed to connect to the scalp compared with 0.1% of novel electrodes. Impedance rose across trials for the conventional cap, whereas the novel attachments showed a smaller increase and maintained lower impedance throughout. Signal quality was significantly better using the novel attachments, while comfort level did not differ between systems. In conclusion, hair-anchored electrode attachment may present a feasible alternative to conventional cap-based EEG for people with Type 4 hair, yielding more usable channels and better signal quality, and supporting more inclusive EEG acquisition.

## 1. Introduction

Electroencephalography (EEG) is widely used across neuroscience, psychology, and clinical research to study brain activity. Yet, Black participants remain underrepresented in EEG studies, limiting the generalizability of findings (Choy et al., 2022; Penner et al., 2023). This gap matters because neurological disorders disproportionately affect Black adults in the United States. Black adults face increased rates of morbidity and mortality from stroke, epilepsy, dementia, and related disorders (Robbins et al., 2022). Stroke risk among middle-aged Black adults, for instance, was nearly three times that of White adults (Howard et al., 2011). Older Black adults experience disproportionate rates of cognitive impairment, with Alzheimer’s disease and related dementias occurring at roughly twice the incidence seen among White adults (Weuve et al., 2018). Closing this gap in EEG research is therefore critical to ensure that EEG measurement is developed with the populations who bear the greatest disease burden.

This underrepresentation in EEG research can be partly attributed to technical limitations of the conventional EEG system. Standard gel-based EEG preparation assumes that hair can be flattened under a cap so that each electrode sits close enough to the scalp to reach an acceptable impedance. However, this assumption does not hold for individuals with Afro-textured hair (Choy et al., 2022; Lees et al., 2024; Webb et al., 2022). Hair geometry varies continuously across the human population, with Afro-textured hair exhibiting the tightest curl patterns among all curl types defined globally (Loussouarn et al., 2007). A commonly used classification describes straight to tightly coiled hair as Types 1-4 (Walker & Wiltz, 1997), with Type 4 further subdivided into 4A-4C by coil pattern (Fig. 1). When a conventional cap is fitted over dense, coily hair, the hair can prevent electrodes from reaching the scalp, increasing electrode-to-scalp distance and compromising contact (Etienne et al., 2020). Even when a recording is obtained, intervening hair can increase noise and reduce signal quality. Consequently, EEG data collected from individuals with coarse and curly hair are rejected at higher rates during quality control (Webb et al., 2022).

**Fig. 1.**
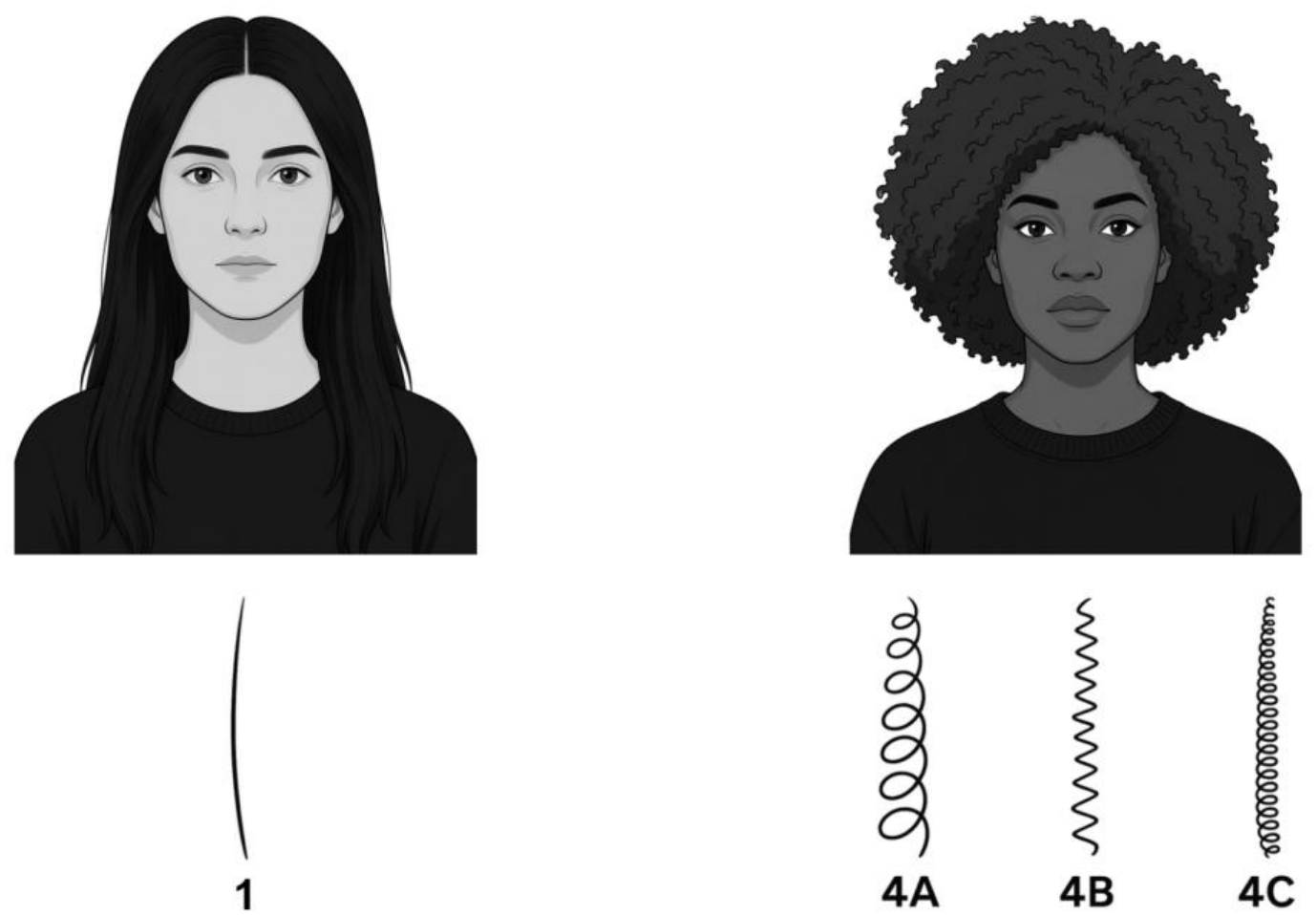
Hair curl contrast between Caucasian/Asian (Type 1) and Black/African American (Type 4). Type 1 hair represents a straight shaft protruding from the hair follicle, while Type 4 hair has a flattened oval shape that then curls upon outer growth. Type 4 hair can be separated into 3 subtypes: 4A (S-shaped coils), 4B (Z-shaped coils), and 4C (tightest coils). Portraits and strand diagrams were generated with artificial intelligence and are approved by the authors for accuracy. They are illustrative and do not depict research participants or research data.

Conventional EEG preparation can impose substantial burdens on participants with Afro-textured hair. Braiding, cornrowing, twisting, and locs are common protective hairstyles among Black individuals (Roseborough & McMichael, 2009), but these hairstyles are frequently treated as incompatible with EEG because scalp contact may require participants to unbraid or alter their hairstyle (Choy et al., 2022). Consequently, Black adults, parents, and children face additional barriers to participating in EEG research because standard protocols ask participants to wear their hair down or alter their hairstyle (Adams et al., 2024). Styling Black hair represents a culturally significant grooming practice that holds deep personal and social importance and serves as a vital protective measure to preserve hair health. Thus, conventional EEG preparation practices can impose disproportionate practical, financial, and cultural burdens on individuals with Afro-textured hair, creating additional barriers to research participation.

Several attempts have been made to overcome the technical challenges to improve the recruitment and retention of Black individuals in EEG studies (Etienne et al., 2020; Kwasa et al., 2024; Richardson et al., 2021). Researchers from BRaIN Lab at the University of Central Florida published a guideline for preparing Afro-textured hair for EEG research to improve signal quality (Richardson et al., 2021). Etienne et al. (2020) developed a Sevo design electrode, which can be placed between cornrows at 10-20 landmarks, resulting in roughly an order of magnitude lower impedance than gold-cup electrodes placed at the same landmarks within braids (Etienne et al., 2020). Kwasa et al. (2024) further evaluated the Sevo design in pediatric epilepsy patients and adults with coarse and curly hair, testing different hair arrangements for electrode placement (unbraided, braided, and Sevo) in clinically relevant outcome measures such as impedance, alpha-band quality, and clinician experience (Kwasa et al., 2024). Together, these studies establish a design principle for hair-inclusive EEG: treat the hairstyle as a structural asset for mounting rather than as a technical obstacle (Etienne et al., 2020; Kwasa et al., 2024).

Building on previous hair-inclusive EEG approaches, we developed 3D-printed attachments that snap onto commercially available wet EEG electrodes and anchor directly within the hair, using the participant’s existing hairstyle, whether loose, braided, or twisted, as the mounting structure rather than relying on a cap pressed over the hair. This study builds on previous work in three complementary ways. First, our study centers on EEG research applications by directly comparing the hair-inclusive approach against the standard cap that most laboratories use (Ricard et al., 2023; Webb et al., 2022). Second, whereas prior work has centered primarily on cornrows, our study aims to accommodate a broader range of natural and protective hairstyles, reducing the burden on participants to restyle their hair before an EEG session (Etienne et al., 2020; Kwasa et al., 2024; Ricard et al., 2023). Third, because hair-inclusive electrodes have, to our knowledge, been evaluated primarily during stationary recordings, we test whether the redesigned attachments remain connected, maintain low impedance, and provide usable EEG signals during walking.

The objective of this study was to design novel EEG electrode attachments and evaluate the technical feasibility and performance of these electrode attachments by comparing them against a commercially available cap system in healthy adults with Afro-textured hair. Our outcome measures include electrode-to-scalp distance, conductive gel usage, electrode disconnection rate, electrode impedance, EEG signal quality, and participant-reported comfort level following both resting and walking trials. We hypothesize that compared with the conventional cap, the novel attachments would reduce electrode-to-scalp distance and conductive gel usage, produce lower electrode disconnections and impedances, improve EEG signal quality, and improve participant-reported comfort following both resting and walking trials. These findings will help determine whether the attachments are feasible for broader research use across Afro-textured hairstyles and will support more equitable adoption of inclusive EEG acquisition methods in neuroscience.

## 2. Methods

### 2.1. Attachment Design

Dense, coily hair increases the distance between the electrode and the scalp and can compromise contact and signal quality. To address this, we designed the attachment to satisfy two requirements: (1) a secure, repeatable fit to a commercial electrode of known dimensions, and (2) sufficient mechanical stability to remain in place during rest and walking.

The attachment was a ring-shaped housing with a snap-fit interface that gripped the electrode directly, modeled in SolidWorks (Dassault Systèmes, France) and Autodesk Fusion 360 (Autodesk, San Francisco, CA, USA), and fabricated on a 3D printer (Anycubic Photon Mono M7, Anycubic, Shenzhen, China) using Anycubic Tough Resin 2.0. We designed three configurations that differ in geometry (the arrangement of prongs that part the hair and let strands pass between them), each intended for a different hairstyle (Fig. 2).

**Fig. 2.**
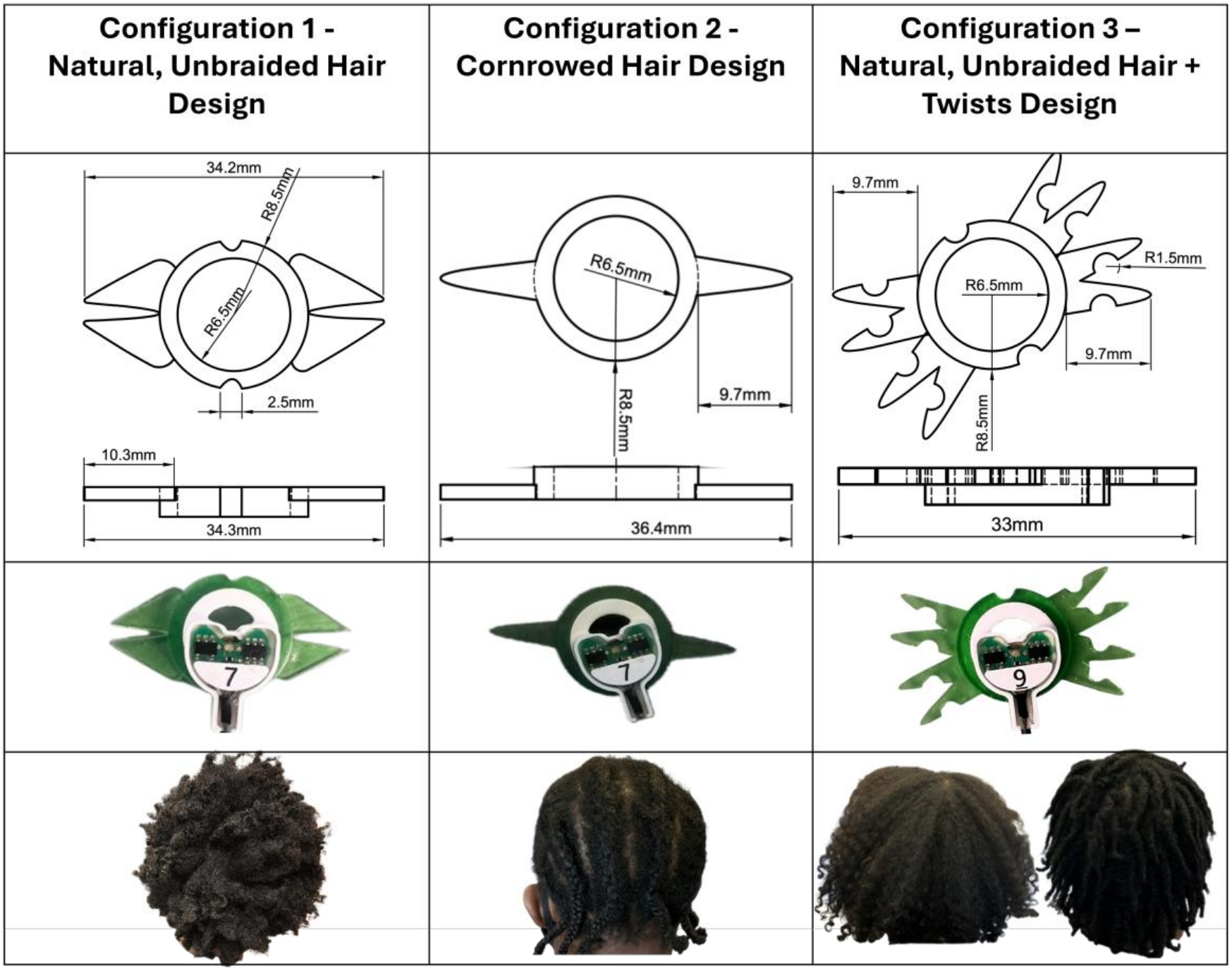
Attachment designs for natural unbraided, braided, twists, and microlocs hairstyles. CAD drawings (top) and 3D-printed configurations (bottom) for the three configurations. All three designs share a common internal ring diameter fitted to the electrode. Representative example hairstyles corresponding to each attachment configuration are shown alongside the designs.

#### Configuration 1 – Natural, Unbraided Hair Design

This design has two thick anchoring prongs, separated by a central slit, and semicircular dents on the central ring (Fig. 2). The dents increase the surface area in contact with loose hair. This lets hair strands pass through the attachment instead of being pushed aside, which keeps the attachment stable against the scalp without braiding. We used this design for unbraided natural Afro-textured hair (subtypes 4b-4c).

#### Configuration 2 – Cornrow Hair Design

This design has two lateral prongs, with prong length matched to braid width (Fig. 2). Prongs were 3D printed in multiple lengths to accommodate different cornrow gaps across participants. We used this design for cornrow hairstyles.

#### Configuration 3 – Natural, Unbraided Hair + Twists Design

This design extends Configuration 1 with eight prongs (four per side), circular dents between adjacent prongs, and matching semi-circular dents on the main ring (Fig. 2). The added prongs and dents increase hair integration and mechanical stability. We designed this configuration for unbraided natural Afro-textured hair (subtypes 4a-4c), twists, and microlocs.

We finalized all three configurations after testing each one directly on mannequin hair and conducting pilot tests to confirm secure anchoring during use. We then assigned each configuration to participants based on their hairstyle during data collection.

### 2.2. Participants

17 healthy adults (12 females, age = 28 ± 10 yr) with Afro-textured hair were recruited from the University of Illinois at Chicago using posted flyers, on-campus outreach, tabling events, and social media announcements. Recruitment started on January 30^th^, 2026, and ended on July 31^st^, 2026. Participants were included if they were 18-60 years old, had Type 4 (4a-4c) hair, self-reported healthy without any significant health conditions (e.g., musculoskeletal, neurological, cardiovascular, or psychiatric), able to walk for 15 minutes with breaks allowed every 3 minutes, and were fluent in English to provide informed consent for participation. One participant was excluded from analysis because the attachment used was not designed for their hairstyle. Among the 16 participants, two participants completed the testing twice, each time with a different novel attachment configuration, yielding 18 entries included in the analyses reported throughout this paper. All participants provided written consent, and the study was approved by the Institutional Review Board at the University of Illinois Chicago (STUDY2025-0685).

We did not perform a formal power calculation because no preliminary data was available to estimate effect sizes, and recruitment was constrained by the one-year timeline of the pilot grant. We targeted five participants per attachment design and enrolled two additional participants for Configuration 3, which was designed to accommodate multiple hairstyles (twists and microlocs).

### 2.3. Experiment Protocol

Participants did not have to alter their hairstyle for Configuration 1 (Natural, unbraided hair design) and 3 (Natural Unbraided Hair + Twists design) attachments and the configuration was selected based on each participant’s hairstyle (Fig. 3). Participants willing to cornrow their hair received written instructions on braid spacing and width and were assigned to Configuration 2 (Cornrow hair design). The instructions made sure the braids fell between electrode sites, so most electrodes could be placed at 10–20 positions without sitting on the cornrows. For Configuration 2, the lateral prong length at each electrode site was chosen to match the cornrow spacing. The most commonly used lengths were 1 cm and 1.4 cm, and 2 cm prongs were used where the cornrows were spaced farther apart.

**Fig. 3.**
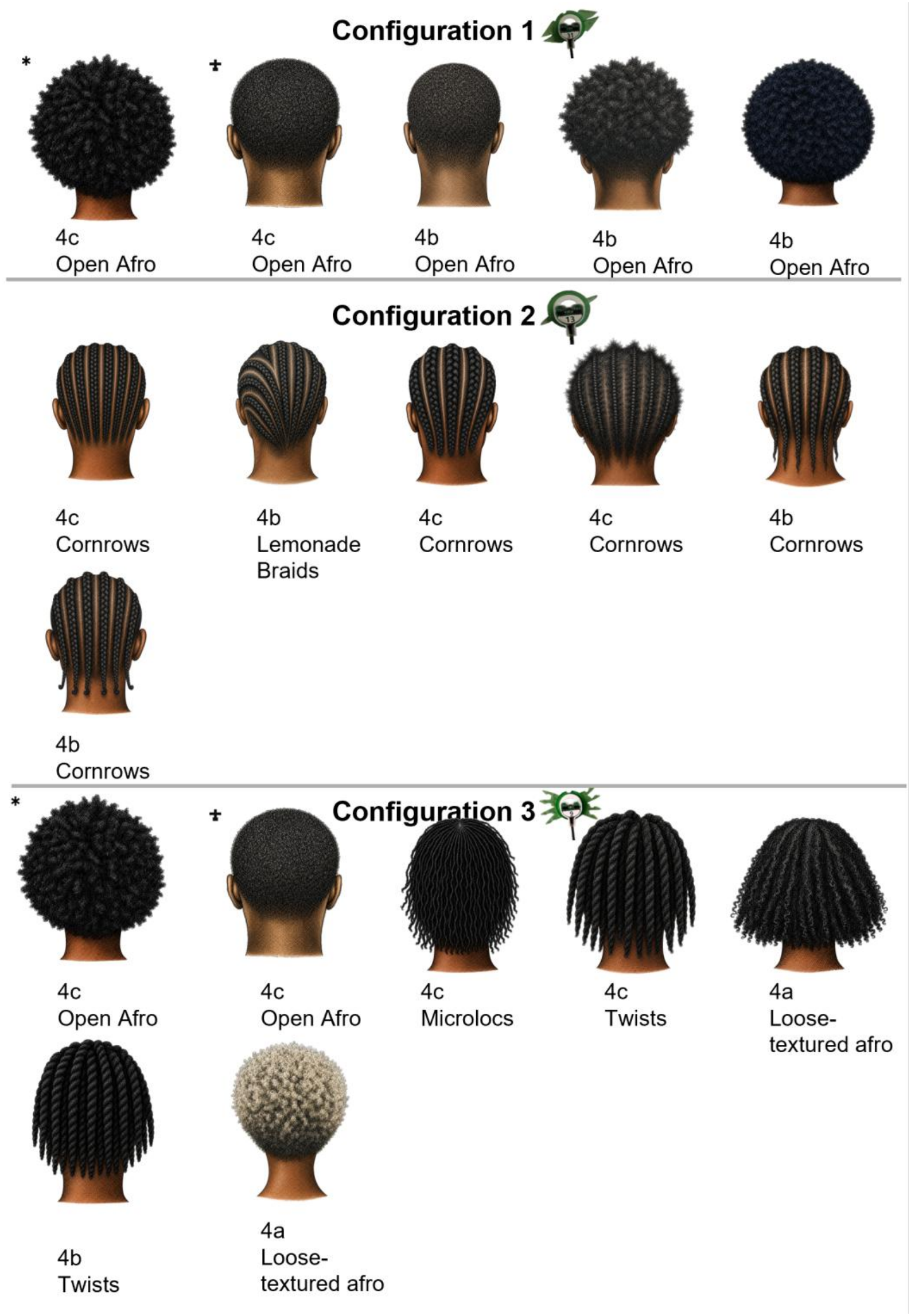
Illustration of hairstyle, hair subtype, hairstyle configurations, and novel attachment assignment for each participant. Hairstyle illustrations were generated with artificial intelligence to represent each participant’s hairstyle at the time of testing. Matching symbols indicate the same participant. Two participants completed the testing twice, each time with a different novel attachment configuration.

EEG was recorded first with a conventional cap (EASYCAP GmbH, Germany) and then with the novel electrode attachments in the single visit. Electrode placement followed the international 10-20 system. For the recording with conventional cap, 16 active electrodes (actiCAP snap electrodes, Brain Products GmbH, Germany) were wired on the cap at left hemisphere sites (Fig. 4A). The ground and the reference were placed at FPz and CPz, respectively. We gelled each electrode with a salt-based conductive gel to make contact with the scalp. We aimed to keep all scalp electrode impedances below 15 kΩ during setup. We added gel until this threshold was reached or until we applied a maximum of 1.5 mL per electrode, whichever came first. We set the 1.5 mL limit based on participants’ self-reported comfort during pilot testing. Impedance was measured using the actiCAP control software (Brain Products GmbH, Germany). We recorded EEG data using LiveAmp 32 amplifiers (Brain Products GmbH, Germany) and logged EEG data at 500 Hz. Participants completed eyes-closed sitting and eyes-open sitting (3 minutes each), standing rest (3 minutes), and walking in the laboratory (2 minutes per trial, 4 minutes total).

**Fig. 4.**
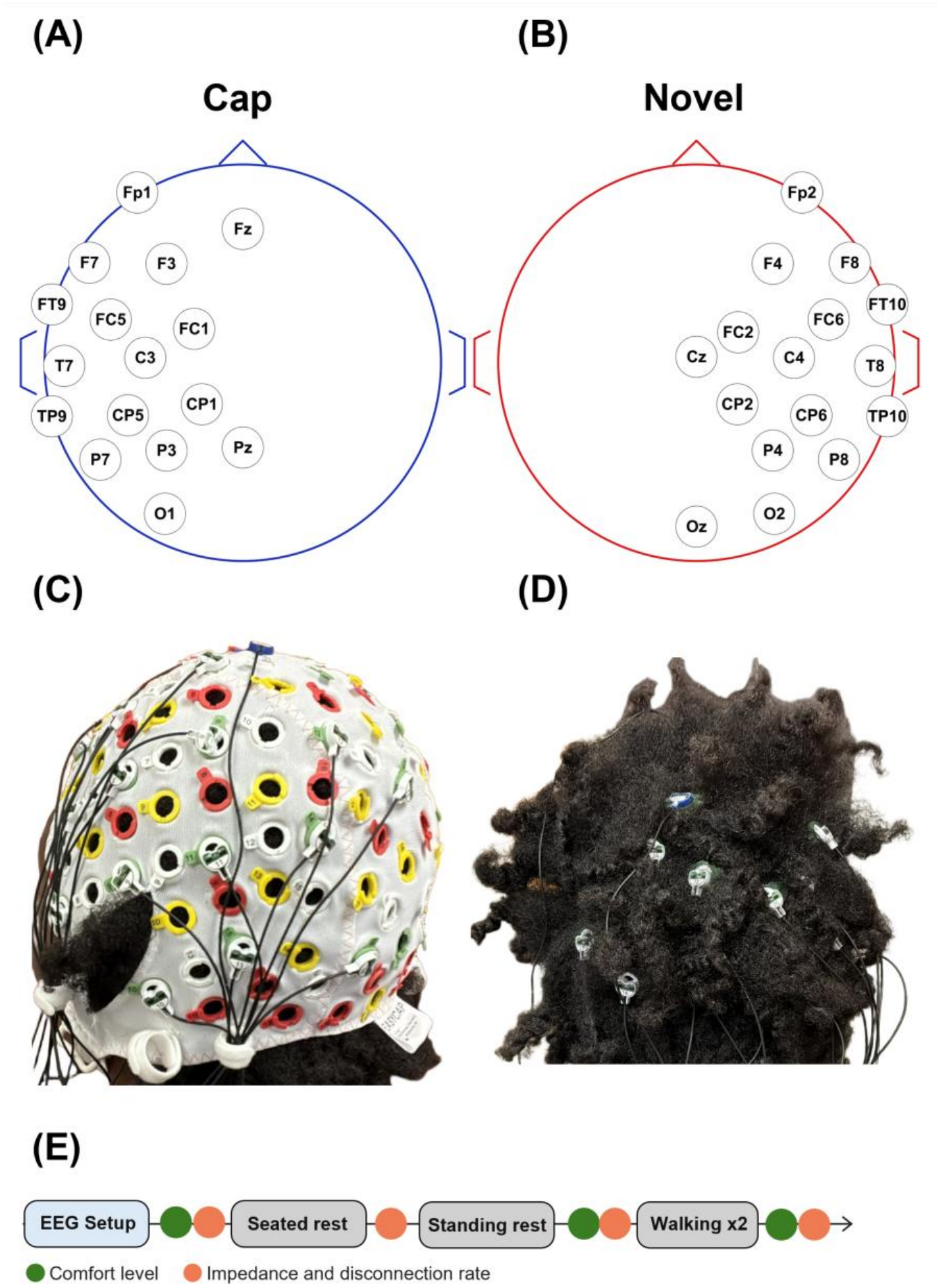
Electrode montages and on-head placement for the conventional cap and novel attachments. (**A**) Conventional cap montage (FP1-O1). (**B**) Novel attachment montage (Oz-FP2). (**C**) Conventional EEG cap fitted on a representative participant with Type 4c hair. We only used 16 electrode slots on the cap. (**D**) Novel 3D-printed attachments placed through the hairstyle on the same participant. (**E**) Experimental session timeline. Green markers indicate when comfort level was taken, and orange markers indicate when impedance and disconnection rate measurements were collected.

The conventional cap was then removed, and a second set of 16 electrodes was placed directly on the scalp at the right hemisphere using the 3D-printed attachments (Fig. 4B). The ground and the reference were placed at the same locations as before. The gelling process was the same as with the conventional cap. Participants then completed the same resting and walking trials. For one participant, the conventional cap was placed on the right hemisphere and the novel attachments on the left to accommodate their hairstyle.

### 2.4. EEG Data Analysis

We performed all analyses in MATLAB R2025b (The MathWorks, Inc., 2026) using the EEGLAB toolbox (Delorme & Makeig, 2004) and custom scripts. We analyzed conventional cap and novel data separately using the electrode positions FP1-O1 and Oz-FP2 respectively (Fig. 4). We applied a 1 Hz high-pass filter with *pop_eegfiltnew (-6 dB at 0.5 Hz)* to remove slow drift, followed by the *CleanLine* to remove 60 Hz line noise. We rejected bad channels that were 3 times the median standard deviation of EEG channels, followed by *pop*_clean_rawdata (flatline criterion = 5 s, channel correlation criterion = 0.8, burst and window criteria disabled), which rejected on average, 2 channels per participant for the conventional cap and 0 for the novel attachments during rest, and 1 and 2 channels per participant during walking. We segmented the resting and walking trial data into non-overlapping 4-second epochs. We computed the power spectral density for each channel using EEGLAB *spectopo (Welch’s method: 2-s Hamming windows, 50% overlap)* and averaged across all 4-s epochs for each trial.

We used the Fitting Oscillations & One Over F (FOOOF) toolbox (version 1.1.1; Donoghue et al., 2020) in Python 3.14.5 to separate the PSDs of the EEG signals into the oscillatory and aperiodic components (1/f) background for each channel (3–40 Hz, fixed aperiodic mode, peak width limits 1–8 Hz, minimum peak height 0.05, maximum 3 peaks). Model goodness of fit was quantified using R². Channel fits were excluded if the estimated aperiodic exponent was negative, as negative exponents are unlikely to reflect physiologically meaningful neural activity (DeVol et al., 2026). We quantified aperiodic-corrected peak alpha power (dB) as 10 times the height of the largest FOOOF-detected oscillatory peak with a center frequency between 8 and 13 Hz above the fitted aperiodic component. For channels in which FOOOF did not identify an alpha peak, aperiodic-corrected peak alpha power was assigned a value of zero. We also retained the aperiodic exponent. We compared only paired conventional cap and novel electrodes at mirrored 10-20 locations, restricting the analysis to electrodes present on both sides of the brain. We used this pairing because more novel electrodes stayed connected than conventional cap electrodes, and pairing kept the number of channels equal between the two systems.

### 2.5. Outcome Measures

Before EEG recording, we collected two preparation-related outcome measures (Fig.4E): electrode-to-scalp distance and gel volume. Electrode-to-scalp distance was measured once per electrode after gelling using a caliper. Gel volume was recorded per electrode from a graduated 10 mL syringe. To track contact stability across the session, we recorded impedance and disconnection rate at four time points: after setup, before standing rest, before walking, and after all trials. An electrode was considered disconnected when no impedance could be measured. At each time point, disconnection rate was calculated as the percentage of disconnected electrodes out of all 16 electrodes.

Additionally, EEG signal quality outcomes were obtained from each trial (eyes-closed sitting, eyes-open sitting, standing rest, and walking) by computing (i) aperiodic-corrected peak alpha power (dB), with higher values indicating stronger alpha signal relative to the background and better signal quality, and (ii) aperiodic component with higher values indicating a steeper slope and less background noise. Lastly, comfort ratings were obtained via a comfort survey scored on a Likert scale (1, not comfortable at all; 10, very comfortable): after setup, after resting trials, and after all trials.

### 2.6. Statistical Analysis

We performed all statistical analyses of the listed outcome measures in R 4.6.1 (R Core Team, 2026) using RStudio (Posit Team, 2026). Significance was set at p < 0.05. Data from the three novel attachment configurations were pooled because configuration was selected according to each participant’s hairstyle and the number of participants using each configuration was insufficient for separate comparisons. Electrode-level measures were averaged within each participant, including electrode-to-scalp distance, gel volume, and impedances.

Preparation-related outcome measures (electrode-to-scalp distance and gel volume) were compared using paired tests. We used the Shapiro-Wilk test to assess normality. If the data were normally distributed, we reported results as mean ± SD and compared with a paired t-test. Effect sizes were reported as Cohen’s d with 0.2, 0.5, and 0.8 indicating small, medium, and large effects, respectively (Cohen, 2009). For non-normally distributed data, results were reported as median [interquartile range, IQR] and compared using Wilcoxon signed-rank tests, reported as Z with effect size r = |Z|/√N, where N is the number of participants. Values of 0.1, 0.3, and 0.5 indicate small, medium, and large effects, respectively (Fiel Peres, 2026).

Electrode disconnection rates were pooled across the four measurement time points, because nearly all novel electrodes remained connected throughout (disconnection rate near 0%). The two systems (conventional cap and novel attachments) were compared with a Wilcoxon signed-rank test. Change in conventional cap disconnection across time points was assessed with a Friedman test (effect size Kendall’s W), followed by pairwise Wilcoxon signed-rank tests with Holm correction.

Impedances were compared between System (conventional cap and novel attachments) and across Time (the four measurement time points) using a two-way repeated-measures ANOVA. Both System and Time were set as within-subject factors. We log-transformed non-normally distributed data.

EEG signal quality outcomes (aperiodic-corrected peak alpha power and the aperiodic exponent) were assessed with a two-way repeated-measures ANOVA separately. System and Condition (eyes-closed sitting, eyes-open sitting, standing rest, and walking) were set as within-subject factors. For the aperiodic exponent, residuals deviated from normality because of extreme cap values from one participant, whose cap had 86% of electrodes disconnected. These values were retained because they reflected genuine recordings of cap performance rather than data errors. A sensitivity analysis excluding this participant did not change the System effect. The System × Condition interaction was sensitive to the sphericity-correction decision, so all data were retained in the primary analysis.

Comfort ratings did not differ across measurement time points for either system (Friedman tests, p ≥ 0.77). Because the primary comparison was overall comfort between conventional cap and novel attachment, we averaged across time points for each participant and compared using the same paired-test procedure as for the preparation-related outcomes.

For all repeated-measures ANOVAs, normality of residuals was assessed with the Shapiro–Wilk test and Q–Q plots. Sphericity was evaluated using Mauchly’s test, and Greenhouse–Geisser-corrected results were reported when the sphericity assumption was violated. Effect sizes are reported as partial η², and significant effects were followed by paired t-tests with Holm correction.

## 3. Results

### 3.1. Electrode-to-scalp Distance

Electrode-to-scalp distance was significantly lower for the novel attachments compared to the conventional cap (t(17) = 8.19, *p* < 0.001, d = 1.93, Fig. 5A). Novel distance was 0.86 ± 0.25 cm, whereas the conventional cap distance was 1.61 ± 0.52 cm. Conventional cap electrodes with the greatest mean distances across participants were CP1 (1.93 cm), Pz (1.90 cm), and O1 (1.79 cm). Novel mean distances remained ≤ 1.0 cm, with the largest distance being Cz (0.98 cm).

**Fig. 5.**
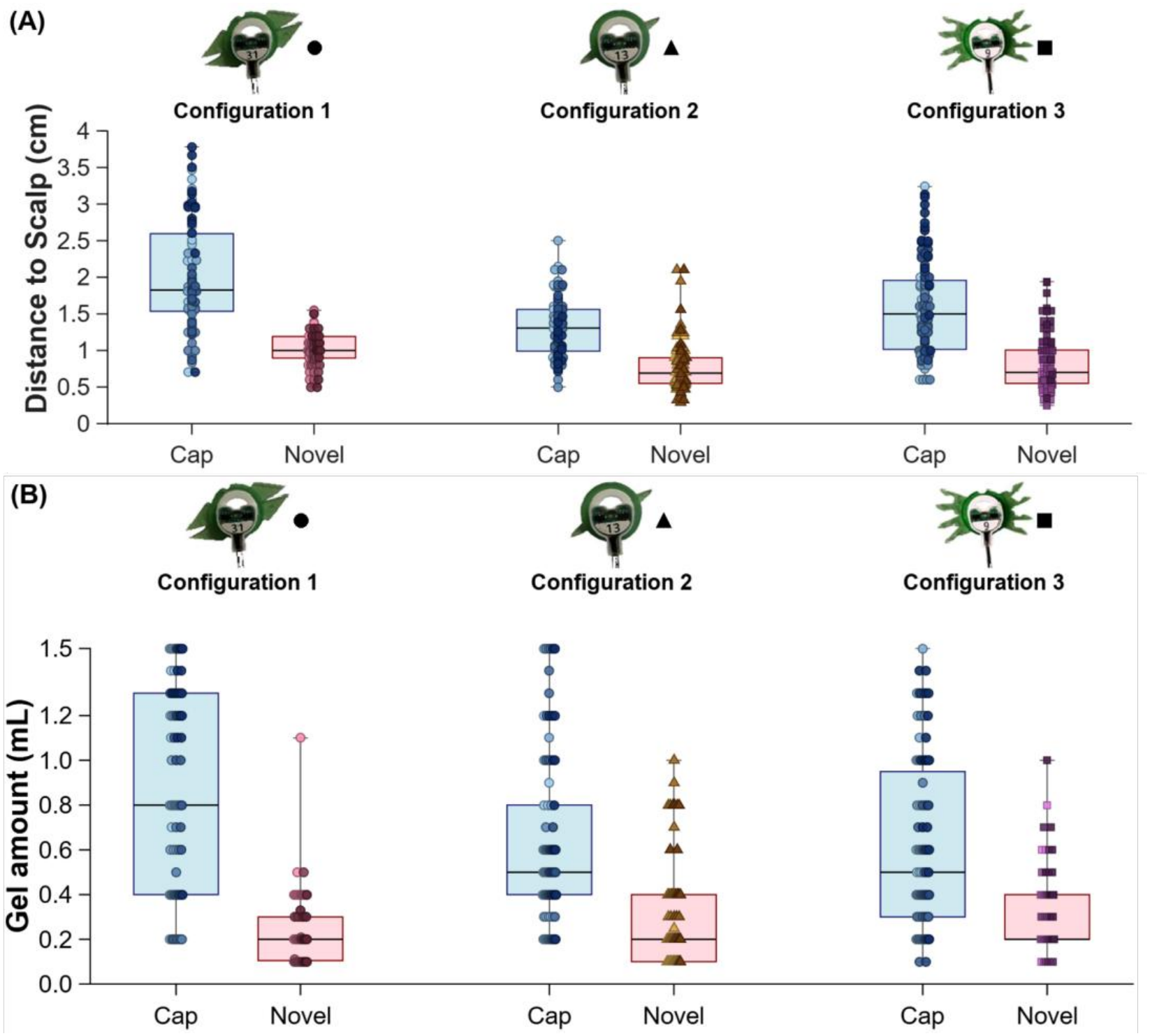
Electrode-to-scalp distance and gel volume box plots (n=18). **(A)** Distance to scalp (cm). **(B)** Gel amount per electrode (mL). Individual participants are shown by novel configuration (Configuration 1-3). Matching marker shade indicates the same participant within each Cap-Novel pair and across each panel. Both distance and gel use were lower for the novel system compared to the cap (p < 0.001).

### 3.2. Gel Usage

Gel usage was significantly lower for the novel attachments compared to the conventional cap (Z = -3.71, *p* < 0.001, r = 0.87, Fig. 5B). Novel gel usage was 0.25 mL (median; IQR 0.20–0.38 mL), whereas the conventional cap gel usage was 0.65 mL (median; IQR 0.52–0.78 mL). Seven of 288 (2.4%) gel readings from the conventional cap electrodes were not recorded as braids or twists prevented syringe access. Conventional cap electrodes with the greatest mean gel use across participants were O1 (0.9 mL) and Pz (0.9 mL). Electrodes with novel attachments remained low, with the highest gel use at O2 (0.4 mL).

### 3.3. Disconnected Electrodes

Electrode disconnection rate was significantly higher for the conventional cap than for the novel attachments (Z = −3.60, p < 0.001, r = 0.85). After setup, the conventional cap had a median of 1 disconnected electrode per participant (range 0–13 of 16), whereas no novel electrodes were disconnected. Across the four measurements, 20.0% of conventional cap electrode readings were disconnected (230 of 1,152; per-participant median 14.8%, IQR 8.2–20.3%), compared with 0.1% for the novel attachments (1 of 1,152; median 0.0%, IQR 0.0– 0.0%). Fig. 6 shows a representative participant’s impedance after all trials, with multiple disconnected conventional cap electrodes and no disconnected novel electrodes.

**Fig. 6.**
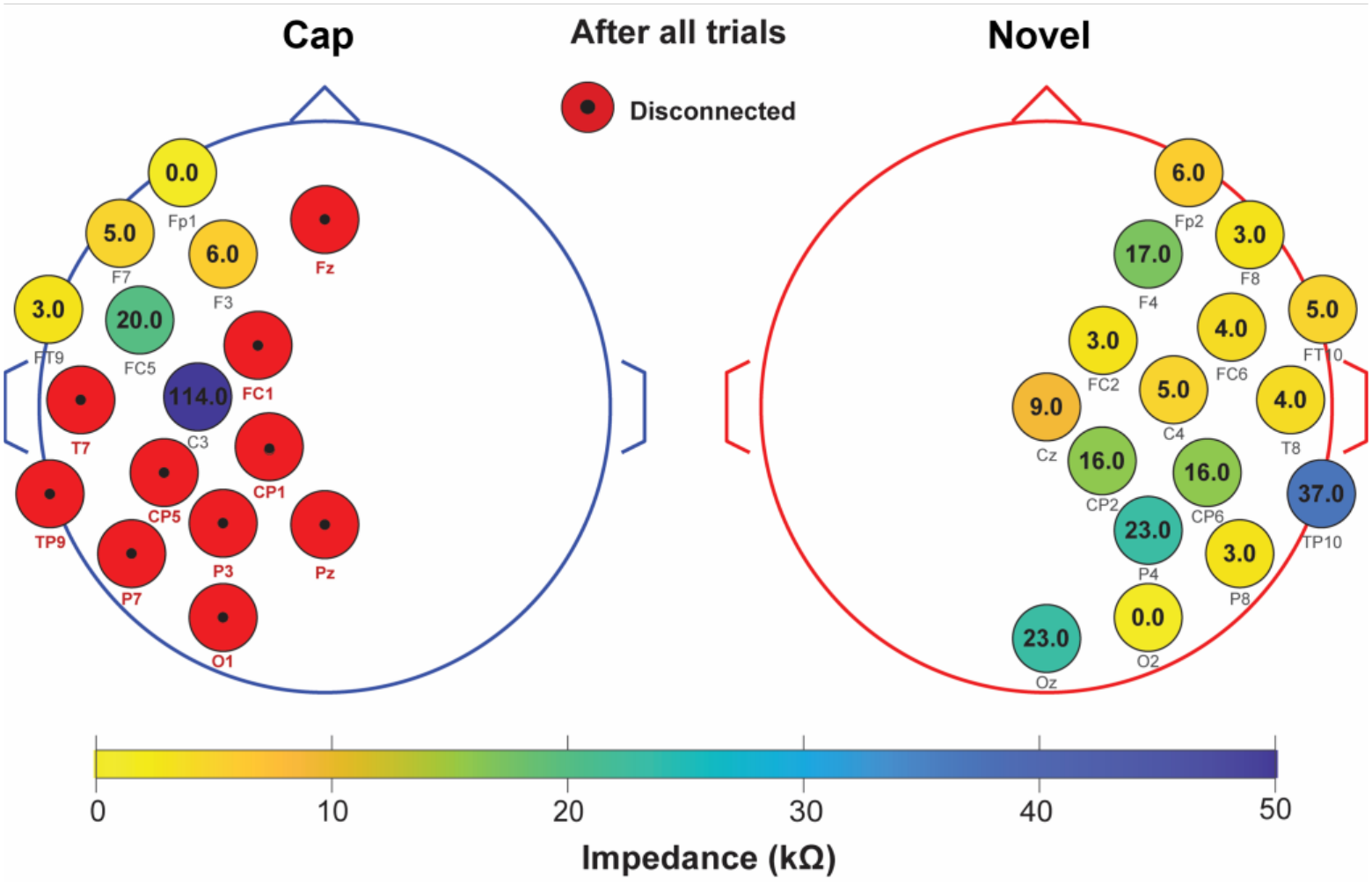
Representative scalp impedance after all trials have concluded for one participant *left – Cap, right - Novel*. Circle color and values represent impedance (kΩ). Disconnected electrodes are marked in red. The conventional cap shows multiple disconnected sites after walking, whereas novel sites remain connected.

Conventional cap disconnection increased across the session (Friedman χ²(3) = 34.92, p < 0.001, W = 0.65), from 12.2% after setup to 19.8% before standing rest, 22.6% before walking, and 25.3% after all trials. Disconnection after setup was lower than at every later time point (p_holm_ ≤ 0.009). The conventional cap electrodes with the highest disconnection rates were TP9 (44.1%), Pz (33.8%), and CP1 and P7 (both 27.9%), whereas the only novel disconnection was a single reading at F8 (Fig. 7).

**Fig. 7.**
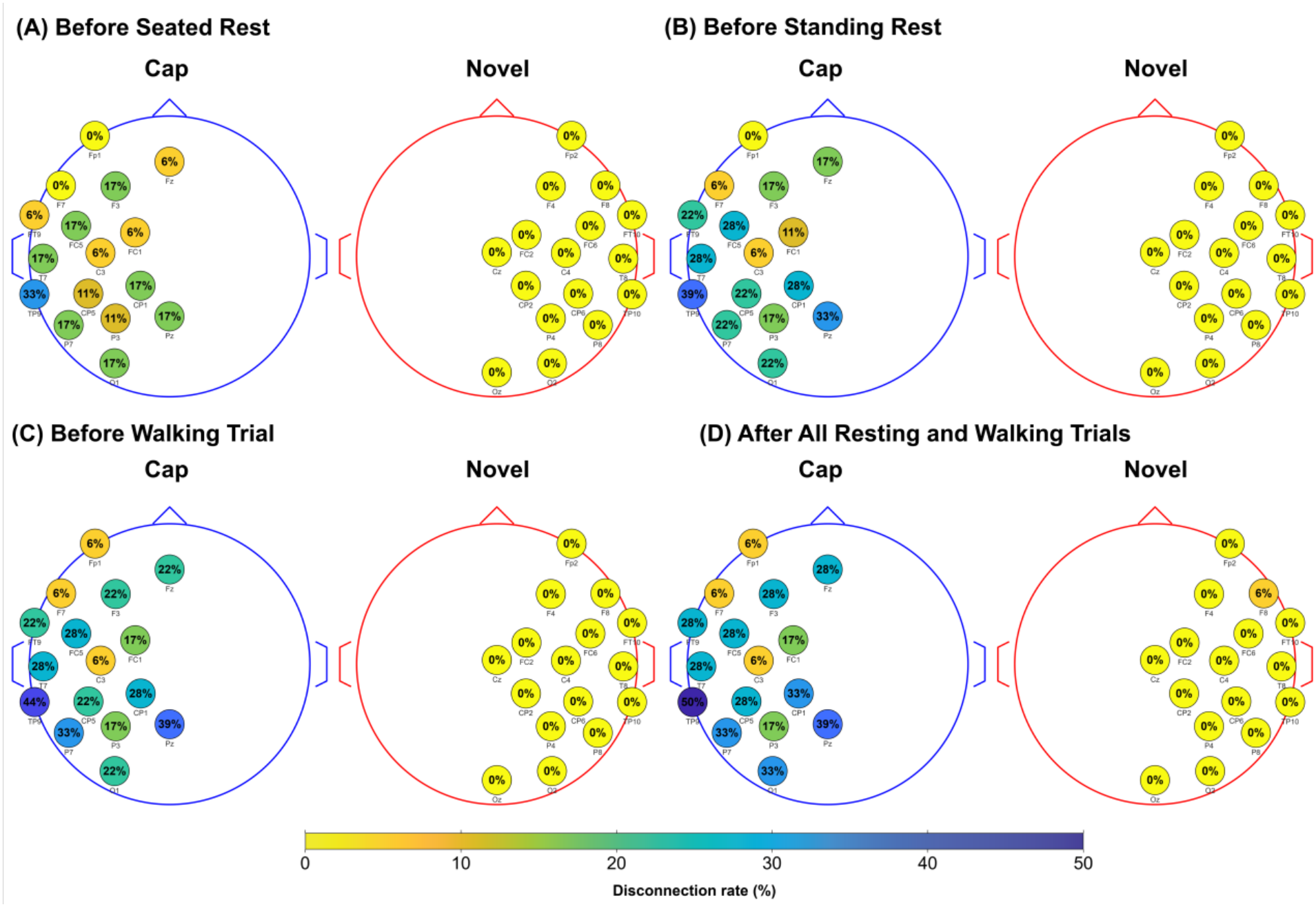
Disconnection rate by electrode location across the cohort (n = 18) at four measurement time points: **(A)** After Setup *left – Cap, right - Novel*, **(B)** Before Standing Rest *left – Cap, right - Novel*, **(C)** Before walking trial *left – Cap, right - Novel*, and **(D)** After all trials *left – Cap, right - Novel*. For each electrode location, and time point, the disconnection rate was calculated as (number of participants with a disconnected reading at that electrode) divided by total number of participants. Circle color represents the percentage of recordings in which that electrode was disconnected at each time point.

### 3.4. Impedance

Using a ≤ 15 kΩ threshold for acceptable impedance, 88.1% (223 of 253) of connected conventional cap electrodes and 98.3% (283 of 288) of novel electrodes achieved low impedance after EEG setup. This decreased to 66.0% (142 of 215) and 91.6% (263 of 287), respectively, after all trials. The conventional cap electrodes with the largest increases from setup were O1 (+26.5 kΩ), FC5 (+19.3 kΩ), and FC1 (+17.8 kΩ), whereas the largest novel increases were at TP10 (+7.2 kΩ) and T8 (+6.2 kΩ), where there was less hair to anchor to (Fig. 8).

**Fig. 8.**
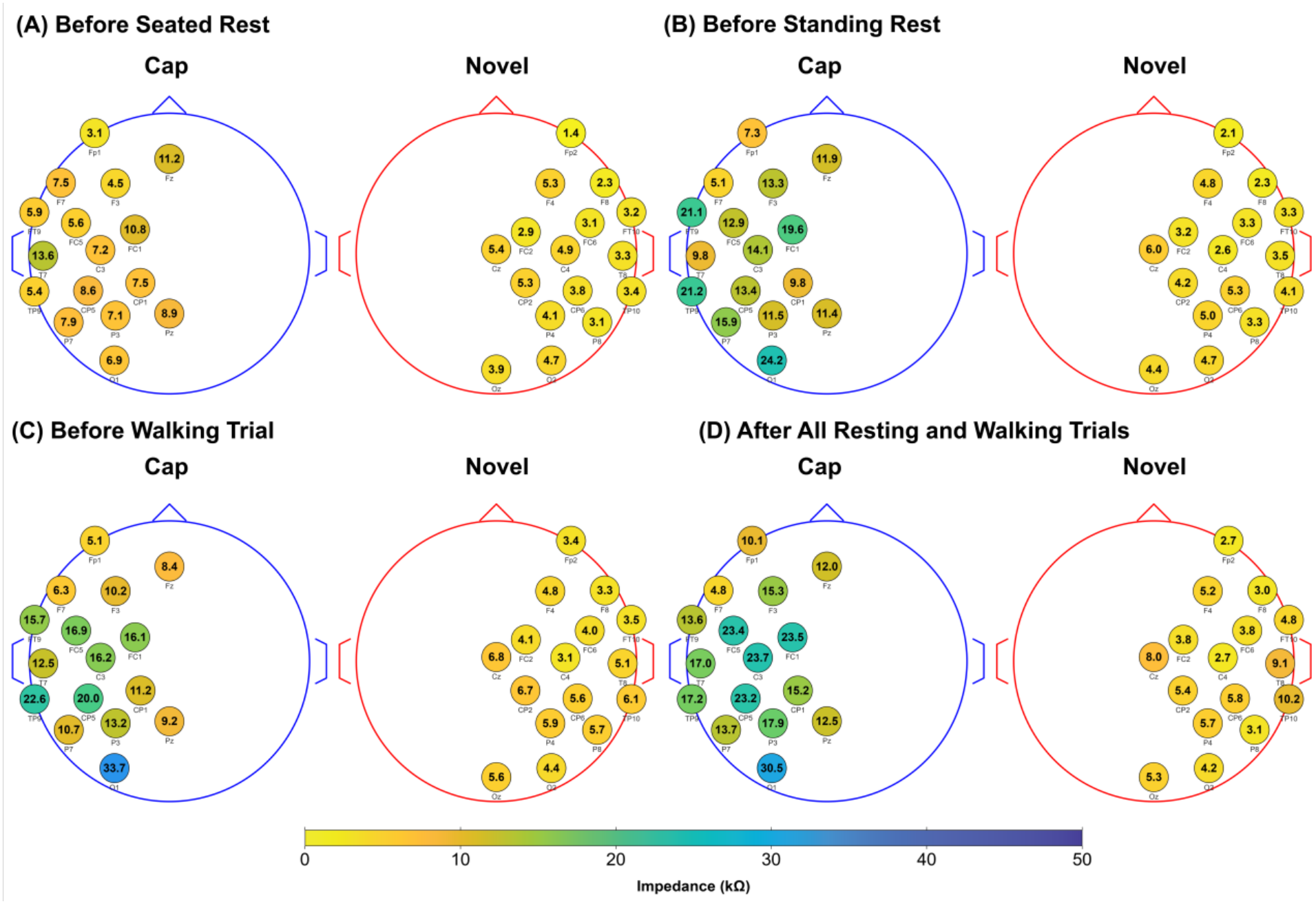
Averaged impedance values (kΩ) across participants (n=18) at four measurement times: **(A)** After setup *left – Cap, right - Novel*, **(B)** Before Standing Rest *left – Cap, right - Novel*, **(C)** Before walking trial *left – Cap, right - Novel*, and **(D)** After all trials *left – Cap, right - Novel*. Circle color indicates the mean impedance at each electrode among connected readings only.

The impedance of connected electrodes was lower for the novel attachments than for the conventional cap, with a significant effect of System (F(1, 17) = 44.66, p < 0.001, partial η² = 0.72; Fig. 9). We also found a significant effect of Time on impedance (F(1.95, 33.07) = 15.89, p < 0.001, partial η² = 0.48) and a significant interaction (F(1.91, 32.48) = 4.12, p = 0.027, partial η² = 0.20), indicating the difference in impedance between systems changed over time. Post hoc comparisons showed that conventional cap impedance was higher than novel impedance at every time point (p_holm_ < 0.01) and rose from 9.0 ± 8.0 kΩ after setup to 17.3 ± 11.0 kΩ after all trials, with all later time points higher than after setup (p_holm_ ≤ 0.010). Novel impedance increased only slightly, from 3.8 ± 1.6 to 5.2 ± 2.4 kΩ, with the only significant increase between before standing rest and after all trials (p_holm_ = 0.009). Lastly, the difference between systems was significantly larger before standing rest than after setup (p_holm_ = 0.018), but not before walking or after all trials (p_holm_ ≥ 0.080).

**Fig. 9.**
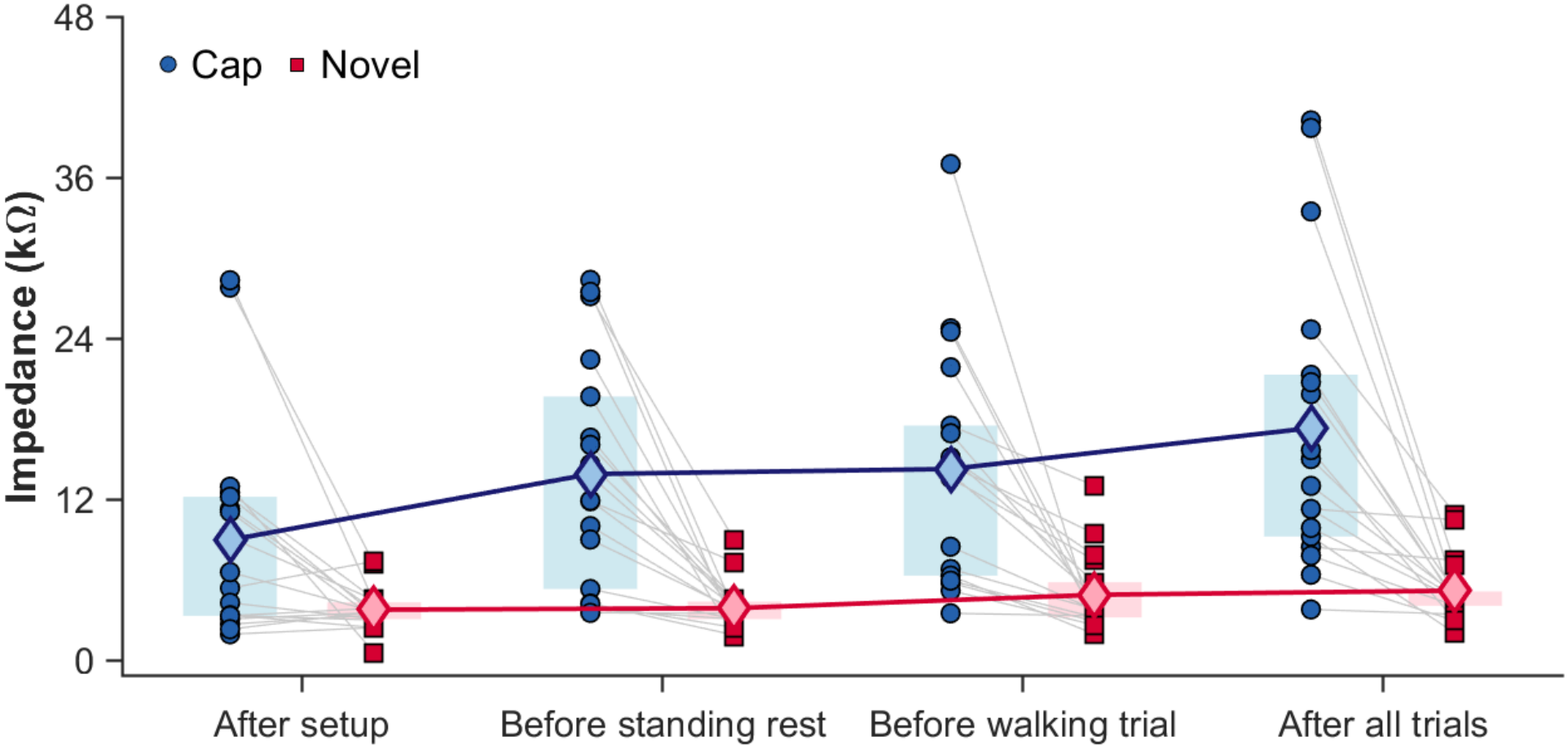
Impedance for the conventional cap and novel attachments after setup, before standing rest, before walking, and after all trials (n = 18). Each circle (cap) or square (novel) represents a participant’s mean impedance across connected electrodes. Shaded regions indicate the interquartile range. Light blue and pink diamonds indicate group means, with colored lines connecting means across time points.

### 3.5. Aperiodic-corrected peak alpha power and aperiodic fit

For the connected electrodes, 10 of 2,714 channel fits (0.4%) were excluded because the aperiodic exponent was negative, and the rest of the channels produced excellent model fit (R² = 0.94 ± 0.10). Of the 2,704 retained channels, aperiodic-corrected peak alpha power was set to 0 dB for 758 channels because FOOOF did not detect a peak between 8 and 13 Hz (211 of 1,597 resting channels and 547 of 1,107 walking channels). Across all resting and walking conditions, conventional cap and novel electrodes were compared at 12 ± 2 pairs (range 5–14).

Aperiodic-corrected peak alpha power was significantly higher for the novel attachments than for the conventional cap (F(1, 17) = 10.35, p = 0.005, partial η² = 0.38; Fig. 10A), suggesting better EEG signal quality with the novel attachments. The interaction between System and Condition was not significant (F(3, 51) = 0.98, p = 0.410, partial η² = 0.05), providing no evidence that the difference between systems depended on condition.

**Fig. 10.**
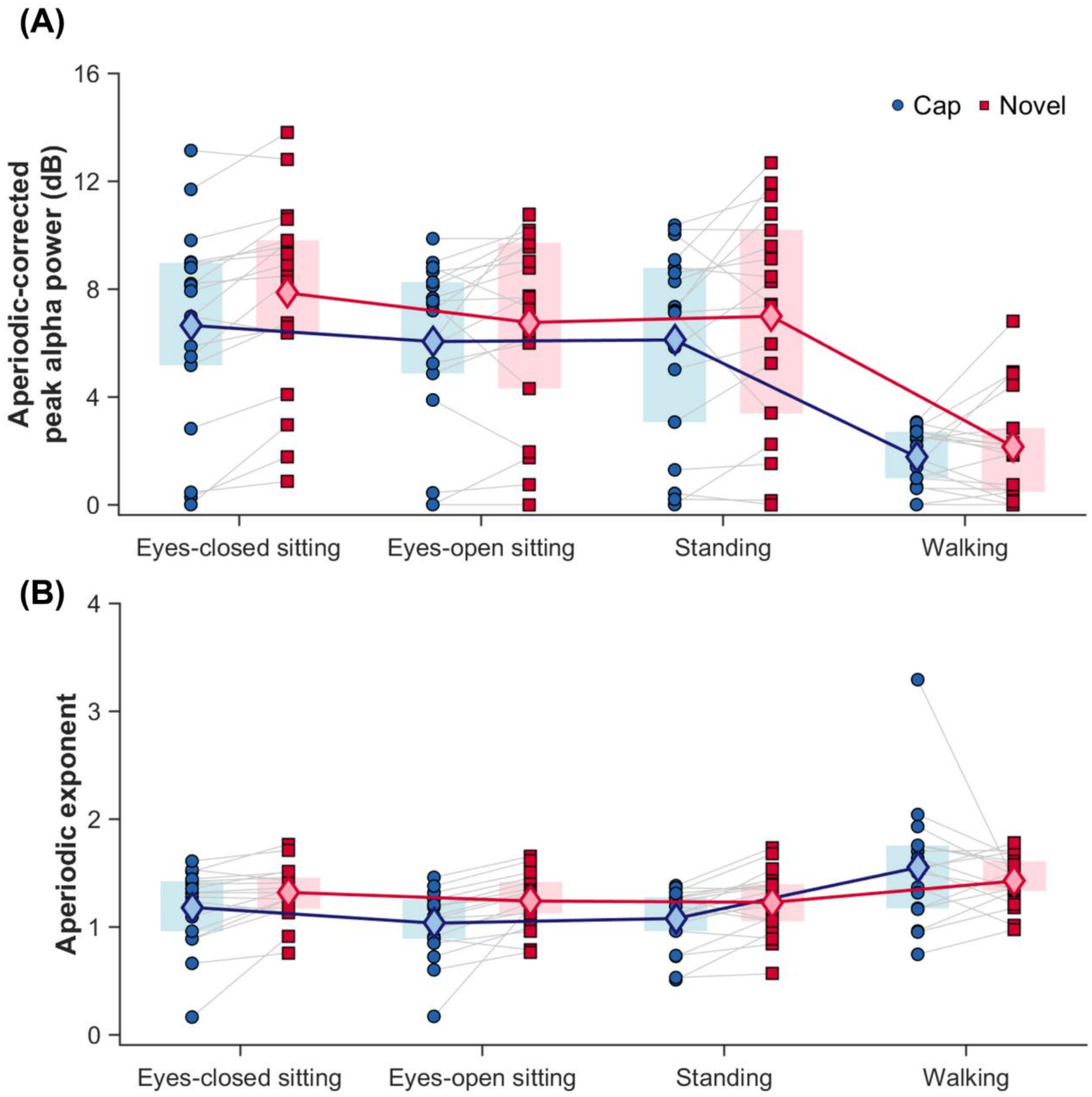
Peak alpha power (A) and aperiodic exponent fit (B) for the conventional cap and novel attachments during eyes-closed sitting, eyes-open sitting, standing rest, and walking (n = 18). Each circle (cap) or square (novel) represents a participant-level value. Shaded regions indicate the interquartile range. Light blue and pink diamonds indicate group means, with colored lines connecting means across conditions.

Aperiodic exponents were significantly steeper for the novel attachments (1.30 ± 0.27) than for the conventional cap (F(1, 17) = 21.06, p < 0.001, partial η² = 0.55; Fig. 10B), suggesting less background noise with the novel attachments. The interaction between System and Condition was not significant (F(1.27, 21.53) = 3.21, p = 0.079, partial η² = 0.16). Excluding the participant with 86% cap disconnection, whose cap values caused the non-normal residuals for the aperiodic exponent, did not change the effect of System (alpha power: p = 0.009; aperiodic exponent: p < 0.001). The interaction for the aperiodic exponent was sensitive to the sphericity-correction decision: the uncorrected result was F(3, 48) = 2.85, p = 0.047, partial η² = 0.15, whereas the Greenhouse–Geisser-corrected result was p = 0.072. Descriptively, the advantage of the novel attachments was smallest during walking, where novel values were higher in only 7 (alpha power) and 9 (aperiodic exponent) of 18 participants.

### 3.6. Comfort Ratings

Comfort ratings did not differ significantly between the conventional cap and novel attachments, although the average comfort level was slightly higher using novel attachments than conventional cap (Z = −1.82, p = 0.069, r = 0.47, Fig. 11). Novel comfort was 9.2 ± 0.9 (median 9.3, IQR 8.8–10.0), whereas conventional cap comfort was 8.4 ± 1.8 (median 9.0, IQR 7.5–9.8). Comfort surveys were introduced after the first three participants had already been enrolled, yielding 15 complete survey responses.

**Fig. 11.**
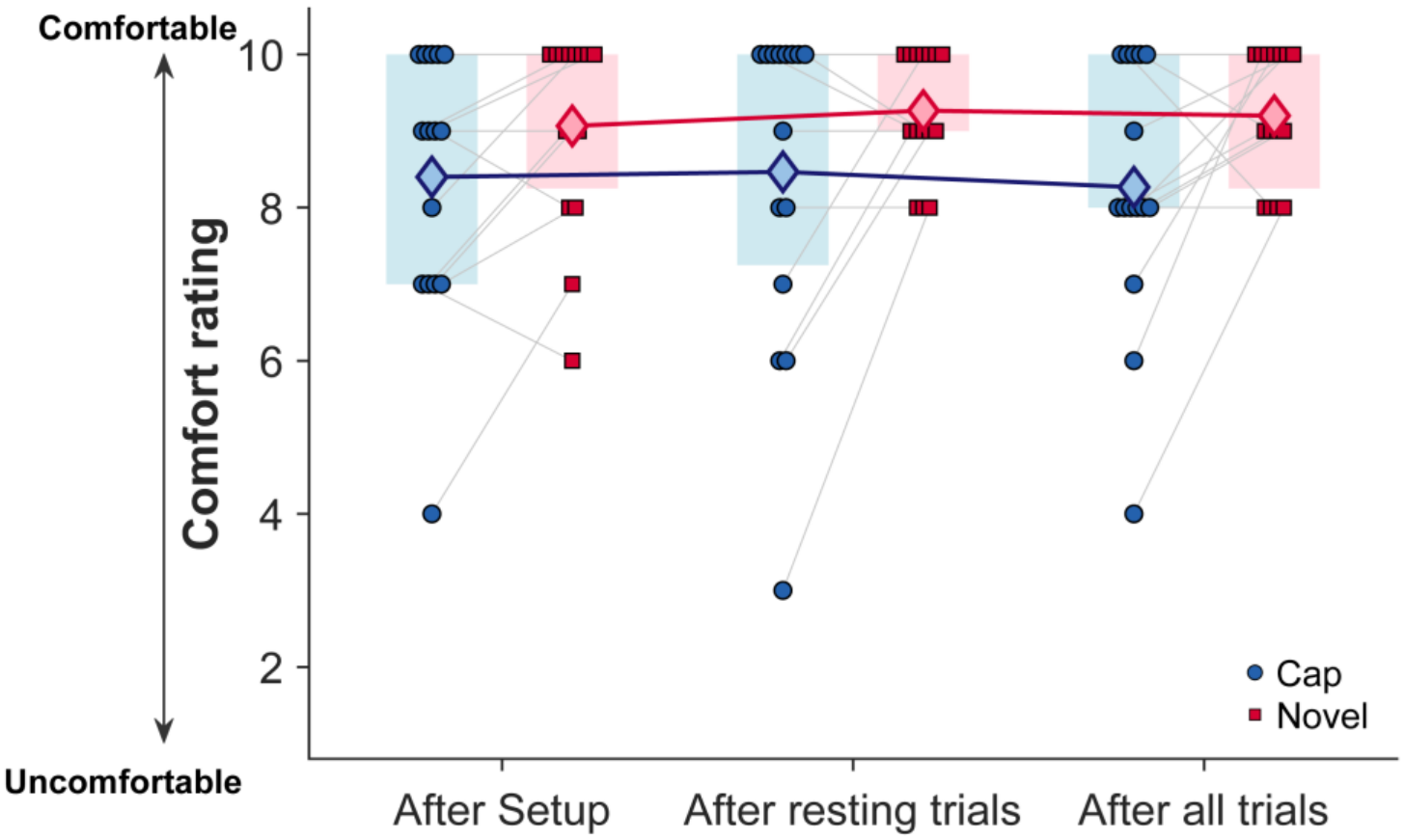
Comfort ratings for the conventional cap and novel attachments after setup, after resting trials, and after all trials (n = 15). Each circle (cap) or square (novel) represents an individual participant’s rating. Shaded regions indicate the interquartile range. Light blue and pink diamonds indicate group means, with colored lines connecting means across conditions. Comfort rating did not differ significantly between the systems.

## 4. Discussion

This study aimed to evaluate the feasibility of novel 3D-printed EEG electrode attachments that secure the electrode directly against the scalp, without a conventional EEG cap, in participants with Afro-textured hair. We designed three electrode attachments to accommodate various hairstyles, including natural loose hair, braids, cornrows, twists, and locs. Compared with the conventional cap, seating the electrodes directly against the scalp reduced the electrode-to-scalp distance, required less conductive gel, lowered the electrode disconnection rate, and kept impedance lower and more stable across the session. EEG signal quality, measured as aperiodic-corrected peak alpha power, was better for novel attachments than conventional cap. These results suggest that hairstyle-anchored attachments are a feasible alternative to low-density cap-based EEG for people with Afro-textured hair, achieving more reliable scalp contact with less gel and without loss of signal quality for the electrodes that remain connected.

### 4.1. Novel attachments present several advantages

The biggest advantage of placing the EEG electrodes with novel attachments directly in the hair, close to the scalp, is that it increases the number of connected electrodes and lowers impedances, enabling EEG collection across all channels. Compared to the conventional cap, the attachments reduced the electrode-to-scalp distance and required a smaller volume of gel, indicating that they seated more directly against the scalp through the hairstyle and therefore needed less gel to establish a connection. That closer seating to the scalp in turn reduced the number of electrodes that disconnected and kept the impedance of connected electrodes lower and more stable across sessions. The reduction in gel matters to participants as well as to data quality, since less residual gel means fewer washes after a session and, plausibly, greater willingness to take part in future EEG research.

Novel electrode attachments enable better connection and low impedance throughout the data collection, whereas conventional cap-based system lost contact progressively. After 9 minutes of resting and 4 minutes of walking trials, only one novel channel had disconnected. The conventional cap showed the opposite pattern as both disconnection rate and impedances increased throughout data collection. Poor initial contact and progressive degradation across rest and walking indicated that conventional cap-based system cannot work well with Afro-textured hair, which left the conventional cap far more prone to complete disconnection while the novel montage remained almost entirely connected. Because disconnected channels are excluded from analysis, reducing those exclusions directly increases usable data.

These results are consistent with prior EEG studies aimed at increasing Black participation that have taken a similar approach. Etienne et al. (2020) and Kwasa et al. (2024) developed Sevo adapters that establish reliable electrode contact with the scalp, reporting lower impedance and favorable participant and technician survey ratings (Etienne et al., 2020; Kwasa et al., 2024). We applied the same principle using 3D-printed attachments that seat the electrode attachment closer to the scalp, with a particular focus on incorporating as many hairstyles as possible. Etienne et al. also reported that Sevo electrode impedances improved relative to their conventional electrodes (Etienne et al., 2020). Our results and prior studies all suggest the feasibility of using a specially designed electrode attachment that leverages the dense, coiled nature of Afro-textured hair to secure EEG electrodes close to the scalp.

Our EEG signal quality was better with novel attachments compared to using the conventional cap. Electrodes with novel attachments showed higher alpha peak power and a steeper aperiodic exponent, indicating that broadband background power fell off faster at higher frequencies than with the conventional cap. Kwasa et al. (2024) similarly proposed a steeper 1/f slope as a signal-quality marker but did not find significant differences between their Sevo design and conventional electrodes (Kwasa et al., 2024). The advantage of novel attachments was clearest at rest. During walking, the novel attachments and cap showed a similar peak alpha power and exponent fit. Therefore, the conventional cap does not perform poorly outright. Using the conventional cap with sufficient connected electrodes may still yield good alpha signal when hair is well prepared and contact is good.

Beyond our present comparison, our novel method for EEG serves as a practical option that other EEG studies can adopt without redesigning their entire recording pipeline. Other research groups can keep their existing EEG hardware and use the 3D-printed attachments for participants with Afro-textured hair. For example, Mirdamadi et al. (2025) showed that a clinically oriented balance N1 measure can be obtained reliably with Cz and two mastoids. In such low-density recordings, replacing those sites with hair-compatible attachments would be a direct approach to increase African American participation and support a more diverse enrollment (Mirdamadi et al., 2025).

### 4.2. Setup time

Setup time is a practical trade-off of the novel attachments. Although a prewired cap can be placed onto the scalp quickly, seating the novel attachments typically took 10 minutes for 16 electrodes, depending on hairstyle. Participants with shorter, open hair, or those with cornrows, took less time for electrode application than those with dense, large afros. Even with this added placement time, however, the novel attachments required less gel to establish contact than the conventional cap, which can help offset the difference in overall preparation time.

### 4.3. Lessons learned from working with Afro-textured hair

We learned some practical lessons that emerged over the course of data collection that may be useful to other groups undertaking hair-inclusive EEG work. Recruitment moved more slowly than we anticipated, particularly during the first half of subject enrollment. We provided an adequate volume of recruitment material, yet many screened individuals did not proceed with enrollment. We also found it easier to recruit participants with open Afro-textured hair compared to the braided configuration. These experiences suggest that other studies incorporating inclusive EEG methods may require community-based education efforts beyond general flyers and pamphlets, especially when the procedure is unfamiliar or requires hairstyle manipulation. Recruitment materials may also benefit from photographs showing Afro-textured hair and explicit language about wash-day timing, so that prospective participants can judge how the study fits their own hair routine before contacting the study team.

The second lesson concerns hairstyle preparation. Some participants were instructed to arrive with a cornrow hairstyle for Configuration 2 attachment. We provided written instructions specifying the exact braid spacing and width the configurations require. Groups adopting this approach should treat hairstyle specifications as part of the protocol, with the same precision as electrode placement, and should anticipate that participant-performed styling will vary.

A third lesson concerns marking electrode locations on darker skin. Light-colored scalp markers (e.g., red or silver) were much easier to see than dark blue and purple, which often blended into dark hair and darker skin and slowed placement. We also found that thicker markers, such as tattoo markers, left a clearer landmark than fine-tipped markers and transferred more reliably from the scalp onto nearby hair strands. Groups collecting EEG with Afro-textured hair should therefore treat marker color and tip width as a crucial part of setup planning and verify visibility before the first participant arrives.

### 4.4. Limitations

Several limitations may constrain the scope of this study and should be considered alongside the main findings. First, we did not counterbalance system order because pressing the conventional cap over hair already gelled for the novel attachments would have smeared gel across the scalp, risking gel bridging between electrodes at the midline. Our preparation-related outcomes and impedances are unlikely to have been affected by order. However, alpha power and the aperiodic exponent can change with time on task and drowsiness, so the signal-quality results should be interpreted with caution. Second, electrodes on the conventional cap were consistently placed on the left hemisphere and the novel electrodes on the right. The hemisphere has little effect on electrode-to-scalp distance, gel volume, disconnection, and impedance, as these are mechanical properties of how an electrode seats in hair. Signal quality, measured via resting alpha power, is normally symmetric in healthy adults. Alpha lateralization has only been reported at frontal sites in psychiatric populations (Avram et al., 2010). Because our participants were screened healthy adults without psychiatric disorders, intrinsic hemispheric differences are an unlikely source of the reported differences between the conventional cap and novel electrode systems. Third, we originally planned to collect electrode displacement measurements using a structural scanner (Structure Sensor 3, model STO3, XRPro) before and after all trials. However, the dense Afro-textured hair constantly obscured the electrodes, preventing reliable measurement. Future studies should use an electrode digitization method that is not image-based.

Although the current attachment housing was designed specifically for actiCAP electrodes, the hair-anchoring prong mechanism is not system-specific and could be adapted to other EEG electrodes by modifying the housing dimensions. Nevertheless, generalizability of the study findings may be limited to a single data collection site and healthy individuals. Participants were recruited at a single university site and prepared by a single team, so results may not generalize to laboratories using different cap sizes, gelling techniques, or levels of experience with Afro-textured hair. We used only 32 channels and evaluated performance only at rest and during walking, leaving higher electrode density and more vigorous movement untested. Participants were also healthy adults, so feasibility in clinical populations, in children, and over the multi-hour recordings typical of clinical monitoring remains to be established.

## Supporting information

STL files for the electrode attachment

## 5. Data Availability

The electrode attachment designs are included in the supplementary material. Code to process the EEG data is available at https://github.com/mbroor/Novel-EEG-Electrodes.

## 6. Acknowledgements

We thank the University of Illinois Chicago Center for Clinical and Translational Science (CCTS) and the Enhancing Cross-disciplinary Innovation and Training Opportunities (EXITO) program in the UIC College of Applied Health Sciences for their support of this work. We also thank the CCTS Community Engagement Advisory Board (CEAB) for their consultation on our study design throughout the project. We also thank Kali Patrice of The Original Hair Whisperer (Personal Care Solutions, LLC) for her guidance on the development of Configuration 3 of our novel attachment design and for her extended support throughout the duration of the study. She met with our team early in the project to discuss hair preparation techniques for highly textured hair relevant to EEG data collection and assisted with participant recruitment. We thank Stanley Donahoo of BrainVision for generously answering our questions about electrode dimensions and for arranging special orders of supplies.

